# Higher classification sensitivity of short metagenomic reads with CLARK-*S*

**DOI:** 10.1101/053462

**Authors:** Rachid Ounit, Stefano Lonardi

## Abstract

The growing number of metagenomic studies in medicine and environmental sciences is creating increasing demands on the computational infrastructure designed to analyze these very large datasets. Often, the construction of ultra-fast and precise taxonomic classifiers can compromise on their sensitivity (i.e., the number of reads correctly classified). Here we introduce CLARK-*S*, a new software tool that can classify short reads with high precision, high sensitivity and high speed at the same time.

## Introduction

One of the primary goals of metagenomic studies is to determine the taxonomical identity of bacteria and viruses in a heterogenous microbial sample (e.g., soil, water, urban environment, human microbiome). This analysis can reveal the presence of unexpected bacteria and viruses in a newly explored microbial habitat (e.g., the marine environment in [1]), or in the case of the human body, elucidate relationships between diseases and imbalances in the microbiome (see, e.g., [2]).

Arguably, the most effective and unbiased method to study these microbial samples is via high-throughput sequencing. The associated computational problem is to assign sequenced (short) reads to a taxonomic unit. While this problem has been studied extensively and several methods and software tools are available, faster and more accurate algorithms are needed to keep pace with the increasing throughput of modern sequencing instruments. In [3] we introduced CLARK, a taxonomy-dependent binning method whose classification speed is currently unmatched. A recent independent evaluation of fourteen taxonomic binning/profiling methods showed that the classification precision of CLARK is comparable (sometimes better) than the state-of-the-art classifiers ([4]). While CLARK’s speed and precision are very high, its classification sensitivity (i.e., the fraction of reads that it correctly classifies) can be significantly improved with the methods described next.

We recall that CLARK is an alignment-free method based on shared *k*-mers. Briefly, it assigns a read *r* to a reference genome *G* if *r* and *G* share more discriminative *k*-mers (i.e., *k*-mers that appear exclusively in one reference genome) than other genomes in the database. Here we show that the classification sensitivity can be increased by allowing mismatches between shared *k*-mers in a limited number of (carefully predetermined) positions, while maintaining the requirement for *k*-mers to be discriminative. The idea of allowing mismatches to improve the sensitivity of seed-and-extend alignment methods was pioneered in [5] with the notion of *spaced seed*. While spaced seeds have been used in some metagenomic binning/profiling methods (e.g., MEGAN [6]), the use of discriminative spaced *k*-mers is novel. Here we describe a major extension of the algorithmic infrastructure of CLARK based on spaced seed, called CLARK-*S.*

## Methods

Given an integer *k* and m reference genomes {*g*_1_, *g*_2_, …, *g*_*m*_}, the set of discriminative *k*-mers *D*_*i*_ for genome *gi* is the set of all *k*-mers in *g*_*i*_ that do not occur (exactly) in any other genome [3]. A spaced seed *s* of length *k* and weight *w* < *k* is a string over the alphabet {1,*} that contains *w* ‘1’ and (*k-w*) ‘*’. Matches are required at a ‘1’ positions, while mismatches are allowed at the ‘*’ locations. The set of discriminative spaced *k*-mers *E_i,s_* is the set of all *k*-mers of *D_i_* that do not occur in any other set *D_j_*(*j* ≠ *i*) when mismatches are allowed at ‘*’ positions in *s*. It is well known that the design of spaced seed is critical to achieve the highest possible precision and sensitivity ([5,7]). Since CLARK is more precise for long contiguous *k*-mers (e.g., *k* = 31), but its highest sensitivity occurs for k in the range [19,22], we considered spaced seeds of length *k*=31 and weight *w*= 22. To determine the optimal positions for the allowed mismatches, we modeled (as it is done in [5]) the succession of ‘1’ and ‘*’ via a Bernoulli distribution with parameter *p*, which represents the similarity level between the read and the genome. We set *p*=0.95 to reflect the expected high similarity between sequences at the species rank. Through an exhaustive search for optimal spaced seeds (with parameters *k* = 31, *w*= 22, *p*= 0.95) using the dynamic programming method by [8] on a region of 100bp, we selected three spaced seeds with the highest hit probability, namely 1111*111*111**1*111**1*11*11111 (hit probability 0.99811), 11111*1**111*1*11*11**111*11111(0.998099), and 11111*1*111**1*11*111**11*11111 (0.998093).

In the preprocessing stage, CLARK-*S* computes and stores on disk, for each genome *g_i_* and each spaced seed *s*, the set of discriminative spaced *k*-mers *E_i,s_*. Compared to the CLARK’s classification phase, CLARK-*S* now requires three look-ups for each *k*-mer in a read (one look-up per spaced seed).

## Experimental Setup

### Database

We compared CLARK-*S* and CLARK on the same set of reference genomes, namely all microbial genomes in the default NCBI/RefSeq database (total of 5,747 species: 1,335 bacteria, 123 archaea and 4,289 viruses).

### Synthetic reads

Evaluations were carried out on simulated datasets and real metagenomic data, as explained next. First, we created six synthetic datasets containing reads from dominant organisms found in the mouth, city parks/medians, gut, indoor and soil environments. A seventh dataset containing reads randomly chosen from 525 bacterial/archaeal species was added (see Supplementary Figures 1-7). These datasets are composed of short synthetic reads generated using ART [9] with default settings (see Supplementary Note 1).

However, observe that a short read *r* generated from genome *g_i_* may appear in another genome for a given error rate or number of mismatches. As a consequence one cannot assume that the “ground truth” of read *r* is *gi*, because *r* might not be unique to *gi*. Ignoring this observation is likely to lead to incorrect conclusions on precision and sensitivity. In order to ensure an unbiased evaluation, we created additional datasets (called “unambiguous”) in which we removed any read that occurs in more than one species, for a given number of allowed mismatches (see Supplementary Note 1 and 2). These datasets only contain <u>unambiguously mapped reads</u> that can allow an unbiased evaluation. In total, we have <u>fourteen datasets containing reads from 647 species</u> (see Supplementary Table 1).

We also added three negative control samples containing short reads that do not exist in any genomes in the NCBI/RefSeq database (see Supplementary Note 1). We used the precision and sensitivity metrics defined [3] to evaluate the classification performance.

### Real metagenomic reads

For experiments on real metagenomes, we chose a large dataset from a recent study on the microbial profile of the NY City subway system, the Gowanus canal and public parks ([10]). We selected twelve samples from various microbial habitat (e.g., bench, garbage can, kiosk, stairway rail, water, etc.), subway stations and riders usage (see Supplementary Table 3). While the ground truth for these data is unknown, the abundance of bacteria, eukaryotes and viruses present in these samples were provided in [10]. Thus, we trimmed raw reads as it was done in [10] (see Supplementary Table 3) and compared the results of CLARK/CLARK-S with the findings in [10] (see Supplementary Table 4 and 5).

## Results

### Synthetic reads

Observe in Supplemental Table 2 that the sensitivity achieved by CLARK-*S* on the fourteen simulated datasets is consistently the highest, while maintaining high precision. Note that the gap in sensitivity is even higher on the unambiguous datasets. On the negative control samples, CLARK-*S* did not classify any reads as expected. Supplemental Table 7 shows that CLARK-*S* classifies about 200 thousand short reads per minute (using one CPU), while CLARK classifies about 3.5 million short reads per minute. If one can take advantage of eight cores, CLARK-S classifies about one million short read per minute, which is sufficiently fast to process large metagenomic datasets in few minutes. CLARK-*S* requires more time to build the database than CLARK, but its RAM usage is comparable (see Supplementary Table 8).

### Real metagenomic reads

Observe in Supplemental Table 6 that CLARK-*S* classifies more reads than CLARK. On average, CLARK-*S* classifies 27% more reads than CLARK. Supplementary Table 5 indicates the reads count assigned by each tool to each species listed in [10] and present in the database. In order to compare results from CLARK/CLARK-S against [10], we estimate the “agreement rate”. For example, in the sample GC01, there are 8 species reported by the study [10] that are present in the database used (i.e., default NCBI/RefSeq genomes of bacteria, archaea and viruses). However, CLARK detected 6 species out of the 8 species, so its agreement rate is 75%. We repeat this estimation for all samples, i.e., for each sample we identified all species detected by [10] that were also present in the database (cf. Supplementary Table 4) and calculate the proportion of species CLARK and CLARK-S detected out of the identified species (cf. Supplementary Table 5).

CLARK-*S* achieves consistently the highest “agreement rate” with [10] on all samples. For instance, in sample P00589 and P00720, CLARK-*S* detected the presence of the virus *Enterobacter phage HK97* but CLARK did not; in sample P01136, CLARK-*S* detected *Brucella ovis* but CLARK did not. In general, CLARK-S identified more relevant organisms than the other tested tools, as observed by a recent study focusing on water samples [11].

## Source code and data

CLARK-S is written in C++ and is freely available at http://clark.cs.ucr.edu. The synthetic datasets (default and unambiguous) are freely available at http://clark.cs.ucr.edu.

## Acknowledgements

This work was supported in part by the US NSF (IIS-1302134 and IIS-1526742).

## Supplementary Material

### Supplementary Note 1: Generation of synthetic datasets and negative controls

In this note, we describe how we created the synthetic datasets used for the evaluation of the three tools we tested. To produce synthetic reads we have considered the species present in real microbial habitats related to mouth, city parks/medians, gut, indoor, and soil (listed below).

- **“Buc12”**: As reported in [4,5], the dominant genus found in the oral cavity is *Streptococcus*. Study [4] also reports the presence of the *Haemophilus influenzae, Haemophilus parainfluenzae, Neisseria subflava* and *Veillonella dispar*. Thus, we selected these four species along with eight species from the *Streptococcus* genus (see Supplementary Figure 1).
- **“CParMed48”**: Forty-eight species were selected from Proteobacteria, Acidobacteria, Bacteroides, Actinobacteria, and *Planctomycetes.* These are the dominant phyla reported in [9] in city parks and medians in Manhattan (see Supplementary Figure 2).
- **“Gut20”**: This dataset contains the twenty species described in the Supplementary Table 1 of [7] (see Supplementary Figure 3).
- **“Hous31”**: Bacteria typically found indoor are *Streptococcaceae, Lactobacillaceae,* and *Pseudomonadaceae* (due to human activities), and also *Intrasporangiaceae* and *Rhodobacteraceae* (due to the environment), as reported in [10] (see Supplementary Figure 4). We selected thirty-one species from these microbial families.
- **“Hous21”**: We selected twenty-one species from the dominant organisms reported in [1] found in the bathroom and kitchen, namely *Propionibacterium acnes, Corynebacterium, Streptococcus,* and *Acinetobacter* (see Supplementary Figure 5).
- **“Soi50”**: We selected fifty species from the dominant genera reported in [3], namely Acidobacteria, Actinobacteria, *Bacteroides, Proteobacteria* and *Verrucomicrobia* (see Supplementary Figure 6).

A seventh dataset “simBA-525” containing reads randomly selected from 525 bacterial/archaeal species was also added (see Supplementary Figure 7). All figures were generated using the Krona tool [8].

#### Datasets generation

We obtained reference genomes from the full NCBI/RefSeq database (∼650 billion of nucleotides, containing more than 58,000 complete genomes distributed in 14,675 species), then we used the ART read simulator [6] to create synthetic reads from the list of species listed above. We ran ART with default quality base profile and error parameters, length 100bp, and coverage 30x. These seven datasets represent a total of 647 species (see Supplementary Table 1 for statistics on these datasets).

#### Unambiguous datasets

To create the “unambiguous” datasets, we used the method described in Supplementary Note 2.

#### Negative control samples

To generate negative controls, we created three datasets (named “LM”, “MH1”, “MH2”) composed of reads that do not exist in any genomes in the NCBI/RefSeq database (see Supplementary Table 1). To build these datasets, observe that if a DNA fragment of 100 bps contains at least one *k*-mer that does not appear in any genomes in the full NCBI/RefSeq database then it does not exist in any of these genomes. In other words, if each read contains one *unassigned k*-mer for the full NCBI/RefSeq database then the read does not map without mismatches (we used *k*=17).

Based on this idea, we generated 10 million 100bp random reads, using a uniform random distribution for each of the four nucleotides (i.e., A, C, G, T have probability 1/4). We also built an index of 17-mers from all genomes in the full NCBI/RefSeq database. Using this index, we counted the number of unknown 17-mers in each random read. Then, we stored one million read that contains at least five unknown 17-mers in dataset “LM”, one million read that contain exactly four unknown 17-mers in dataset “MH1”, and one million read that contain exactly three unknown 17-mers in dataset “MH2”.

### Supplementary Note 2: Generation of “unambiguous” datasets

In this note, we describe how to create datasets with unambiguously mapped reads, from the set of reads generated by ART.

#### Definitions and notations

Given a string *x*, let *|x|* denote its length.

<u>Definitions</u>: In the following definition we assume that *k* is a positive integer (length of the k-mers), *r* is a read, and *G* is a genome.

- Given a set of genomes {*G*_1_, *G*_2_, …, *G*_*m*_}, a *k*-mer *T* is *specific* to *G_i_* if *T* occurs in *G_i_* (exactly) but *T* does not occur (exactly) in any other genome *G_j_*, when *j ≠ i*.
- Given a set *K* of *k*-mers specific to *G*, the number of nucleotides of read *r* covered by at least one *k*-mer in *K* is called the *coverage* of *r* to *G* which we denote by *cov(r,G)*.
- Given a position *l* ∈ *[1,|G|-|r|+1],* we denote by *M(r,G,l)* the number of mismatches (Hamming distance) between read *r* and a substring of *G* of length *|r|* starting at position *l*.
- We denote by *OPT(r,G)* = min_*l* ∈ *[1,|G|-|r|+1]*_ *{M(r,G,l)},* i.e., the minimum number of mismatches for all possible alignments (with no indels) between *r* and *G*.
- Given a set of genomes {*G*_1_, *G*_2_, …, *G_m_*}, read *r* is *unambiguously mapped* to *G*_*i*_ if and only if for all *j ≠ i* we have that *OPT(r,G_i_) < OPT(r,G_j_)*. In other words, there is no pair of genomes (*G_i_*, *G_j_*) such that the two optimal alignments of *r* to *G_i_* and *G_j_* achieves the same number of mismatches.

<u>Lemma</u>: Given a read *r*, a positive integer *k* and a set of genomes {*G*_1_, *G*_2_, …, *G*_*m*_} if there exists an index *i* ∈ *[1, m]* such that if ⌊*cov(r, G_i_)/k⌋*> *M(r,G_i_)* then for all *j* ≠ *i*, we have that *OPT(r,G_j_)* > *OPT(r,G_i_)*.

*Proof*: By definition of *k*-mer specific to a genome: for each non-overlapping block *B* of *k* nucleotides that are covered by at least one *k*-mer specific to *G_i_* in *r*, at least one mismatch exists between *B* and any block of *k* nucleotides in *G_j_* where *i* ≠ *j*. Since there is at least ⌊*cov(r,G_i_)/k*⌋ non-overlapping block(s) of *k* nucleotides covered by at least one *k*-mer *G_i_*-specific in *r*, for all *j* ≠ *i* we have that *OPT(r,G_j_)* ≥ ⌊*cov(r,G_i_)/k*⌋. By definition, we have that *OPT(r,G_i_) ≤ M(r,G_i_). For all *j* ≠ *i*, *OPT(r,G_j_)* ≥ ⌊cov(r,G_i_/k*⌋ and, by the hypothesis of the lemma, we have that ⌊*cov(r,G_i_)/k*⌋ > *M(r,G_i_)* implies that *OPT(r,G_j_)* ≥ ⌊*cov(r,G_i_)/k*⌋ > *M(r,G_i_)* ≥ *OPT(r,G_i_)*. Thus, for all *j* ≠ *i*, *OPT(r,G_j_)* > *OPT(r,G_i_)*.

In other words, if ⌊*cov(r,G_i_)/k*⌋ is higher than the number of mismatches between *r* and *G_i_* then the read r is unambiguously mapped to G_i_.

<u>Unambiguously mapped reads</u>: We used the ART read simulator to create simulated datasets. We considered the species rank, so genomes of the same species were considered together as a unique sequence. We set *k*=19 to determine sets of *k*-mers specific to each species (i.e., 14,675 sets), then we created a hash-table to extract all 19-mers from all species and remove all 19-mers that are common to at least one pair of species. To create a dataset of unambiguously mapped reads, we filtered reads as follows. For each species *G* of a given dataset, and for each read *r* created, we use the alignment (provided by ART) of *r* to its reference sequence of origin. We compute the number of mismatches *M* between *r* and *G*, and we estimated the specificity-coverage *C* of *r* to *G*. Using the previous Lemma, *r* was added to the unambiguous variant of the dataset (because it is unambiguously mapped to *G*) if the value *C/k* was higher than *M+* 1.

**Supplementary Table 1:** Number of reads and species in each synthetic datasets (default and unambiguous) and for the negative controls.

| Synthetic datasets | *Buc12* | *CParMed48* | *Gut20* | *Hou31* | *Hou21* | *Soi50* | *simBA-525* |
| --- | --- | --- | --- | --- | --- | --- | --- |
| Species | 12 | 48 | 20 | 31 | 21 | 50 | 525 |
| Reads (default) | 600,000 | 1,200,000 | 500,000 | 775,000 | 525,000 | 2,500,000 | 5,666,143 |
| Reads (unambiguous) | 600,000 | 1,200,000 | 500,000 | 750,000 | 500,000 | 2,500,000 | 5,727,654 |

| Negative control | *HM1* | *HM2* | *LM* |
| --- | --- | --- | --- |
| Reads | 1,000,000 | 1,000, 000 | 1,000,000 |

**Supplementary Table 2:**
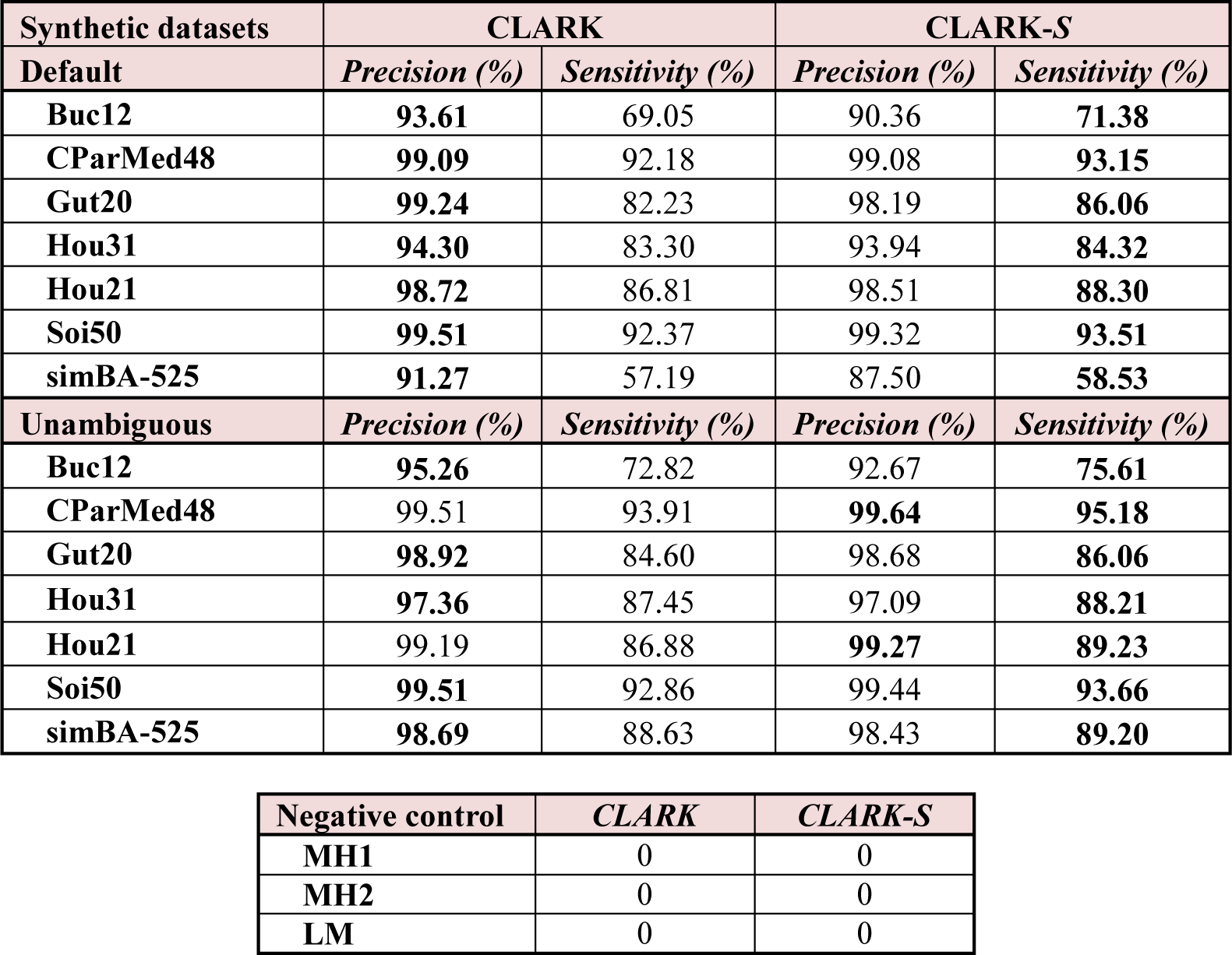
Precision and sensitivity for CLARK, and CLARK-*S* on the synthetic datasets (default, unambiguous). The highest value for precision and sensitivity are indicated in bold. The second table reports the count of classified reads for CLARK and CLARK-*S* for the negative controls.

| Synthetic datasets | CLARK |  | CLARK-S |  |
| --- | --- | --- | --- | --- |
| Default | *Precision (%)* | *Sensitivity (%)* | *Precision (%)* | *Sensitivity (%)* |
| Buc12 | **93.61** | 69.05 | 90.36 | **71.38** |
| CParMed48 | **99.09** | 92.18 | 99.08 | **93.15** |
| Gut20 | **99.24** | 82.23 | 98.19 | **86.06** |
| Hou31 | **94.30** | 83.30 | 93.94 | **84.32** |
| Hou21 | **98.72** | 86.81 | 98.51 | **88.30** |
| Soi50 | **99.51** | 92.37 | 99.32 | **93.51** |
| simBA-525 | **91.27** | 57.19 | 87.50 | **58.53** |
| Unambiguous | *Precision (%)* | *Sensitivity (%)* | *Precision (%)* | *Sensitivity (%)* |
| Buc12 | **95.26** | 72.82 | 92.67 | **75.61** |
| CParMed48 | 99.51 | 93.91 | **99.64** | **95.18** |
| Gut20 | **98.92** | 84.60 | 98.68 | **86.06** |
| Hou31 | **97.36** | 87.45 | 97.09 | **88.21** |
| Hou21 | 99.19 | 86.88 | **99.27** | **89.23** |
| Soi50 | **99.51** | 92.86 | 99.44 | **93.66** |
| simBA-525 | **98.69** | 88.63 | 98.43 | **89.20** |

| Negative control | *CLARK* | *CLARK-S* |
| --- | --- | --- |
| MH1 | 0 | 0 |
| MH2 | 0 | 0 |
| LM | 0 | 0 |

**Supplementary Table 3:** Metadata of the selected real samples from [2]: Sample ID, number of raw reads, number of reads after trimming, object swabbed, location of the sample, borough name, and the number of weekly riders in 2013. Raw reads were trimmed as done in [2]: the first/last 10bp each read were removed (reads longer than 100bp were truncated and the first 100bp were kept); trimmed reads with more than 10 bases with quality scores less than 20 were removed.

| **Sample ID** | **Raw reads** | **Trimmed reads** | **Object swabbed** | **Location** | **Borough** | **Weekly riders** |
| --- | --- | --- | --- | --- | --- | --- |
| **GC01** | 29,282,945 | 28,739,916 | Water Sample | Gowanus Canal | Brooklyn | NA |
| **P00090** | 3,161,196 | 3,085,871 | Stairway rail | Times Sq-42 St/42 St | Manhattan | 197,696 |
| **P00302** | 12,206,080 | 11,700,388 | Bench | 59 St-Columbus Circle | Manhattan | 72,236 |
| **P00306** | 7,536,640 | 7,194,993 | Kiosk | 34 St-Penn Station | Manhattan | 90,042 |
| **P00454** | 7,872,512 | 7,555,783 | Bench | Fulton St | Manhattan | 64,461 |
| **P00589** | 3,129,344 | 3,015,949 | Turnstile | Broadway-Lafayette St/Bleecker St | Manhattan | 38,799 |
| **P00720** | 6,833,000 | 6,536,830 | Bench | Franklin St | Manhattan | 5,825 |
| **P00945** | 7,530,914 | 7,257,415 | Bench | Forest Av | Queens | 4,103 |
| **P01041** | 1,171,456 | 1,160,282 | Bench | Van Siclen Av | Brooklyn | 2,974 |
| **P01136** | 6,417,114 | 6,220,889 | Garbage Can | Jefferson St | Brooklyn | 6,612 |
| **P01270** | 17,072,185 | 16,471,331 | Seats | F Train | Brooklyn | NA |
| **P01324** | 2,686,976 | 2,594,672 | Garbage Can | Whitlock Av | Bronx | 1,685 |

**Supplementary Table 4:** List of species detected in [2] which are also present in the database (i.e., bacteria/archaea/viruses genomes from NCBI/RefSeq) for each of the twelve samples.

| Sample ID | Species in [2] and present in the default RefSeq database (bacteria/archaea/viruses) |
| --- | --- |
| GC01 | *Bifidobacterium adolescentis*, *Bifidobacterium longum*, *Desulfobacterium autotrophicum*, *Erwinia billingiae*, *Eubacterium eligens*, *Eubacterium rectale*, *Methanocorpusculum labreanum*, *Parabacteroides distasonis* |
| P00090 | *Acinetobacter baumannii*, *Cronobacter turicensis*, *Enterobacter cloacae*, *Enterococcus casseliflavus*, *Enterococcus faecalis*, *Klebsiella pneumoniae*, *Lysinibacillus sphaericus*, *Macrococcus caseolyticus*, *Micrococcus luteus*, *Pseudomonas putida*, *Pseudomonas stutzeri*, *Stenotrophomonas maltophilia*, *Streptococcus suis* |
| P00302 | *Achromobacter xylosoxidans*, *Acinetobacter baumannii*, *Bacillus megaterium*, *Dickeya dadantii*, *Enterobacter cloacae*, *Enterococcus casseliflavus*, *Enterococcus faecalis*, *Enterococcus faecium*, *Enterococcus hirae*, *Fingoldia magna*, *Klebsiella pneumoniae*, *Lactococcus lactis*, *Leuconostoc mesenteroides*, *Lysinibacillus sphaericus*, *Micrococcus luteus*, *Propionibacterium acidipropionici*, *Propionibacterium acnes*, *Pseudomonas putida*, *Pseudomonas stutzeri*, *Staphylococcus epidermidis*, *Staphylococcus haemolyticus*, *Stenotrophomonas maltophili* |
| P00306 | *Acinetobacter baumannii*, *Acinetobacter oleivorans*, *Enterobacter cloacae*, *Enterobacteria phage IME10*, *Enterococcus casseliflavus*, *Enterococcus faecium*, *Klebsiella pneumoniae*, *Propionibacterium acnes*, *Pseudomonas stutzeri*, *Stenotrophomonas maltophilia* |
| P00454 | *Acinetobacter baumannii*, *Chlorobium phaeobacteroides*, *Enterobacter cloacae*, *Enterococcus casseliflavus*, *Enterococcus mundtii*, *Klebsiella pneumoniae*, *Lysinibacillus sphaericus*, *Pseudomonas stutzeri*, *Solibacillus silvestris*, *Stenotrophomonas maltophilia* |
| P00589 | *Acinetobacter baumannii*, *Enterobacter cloacae*, *Enterobacteria phage HK97*, *Enterococcus casseliflavus*, *Lactococcus lactis*, *Pseudomonas putida*, *Pseudomonas stutzeri*, *Streptococcus suis* |
| P00720 | *Corynebacterium variabile*, *Enterobacter cloacae*, *Enterobacteria phage HK97*, *Enterococcus casseliflavus*, *Lactococcus lactis*, *Leuconostoc citreum*, *Lysinibacillus sphaericus*, *Pseudomonas stutzeri*, *Stenotrophomonas maltophilia* |
| P00945 | *Bacillus megaterium*, *Enterobacter cloacae*, *Enterococcus faecalis*, *Enterococcus faecium*, *Lysinibacillus sphaericus*, *Pseudomonas putida*, *Pseudomonas stutzeri*, *Stenotrophomonas maltophilia*, *Stenotrophomonas phage phiSMA7* |
| P01041 | *Enterobacter cloacae*, *Enterobacteria phage HK97*, *Enterococcus casseliflavus*, *Enterococcus faecalis*, *Pseudomonas stutzeri*, *Stenotrophomonas maltophilia* |
| P01136 | *Brucella ovis*, *Corynebacterium variabile*, *Enterobacter cloacae*, *Enterobacteria phage HK97*, *Enterococcus casseliflavus*, *Leuconostoc mesenteroides*, *Pseudomonas putida*, *Pseudomonas stutzeri*, *Stenotrophomonas maltophilia*, *Streptococcus suis* |
| P01270 | *Achromobacter xylosoxidans*, *Enterobacter cloacae*, *Enterococcus casseliflavus*, *Enterococcus faecalis*, *Enterococcus faecium*, *Enterococcus hirae*, *Lactococcus lactis*, *Lysinibacillus sphaericus*, *Propionibacterium acnes*, *Pseudomonas putida*, *Pseudomonas stutzeri*, *Stenotrophomonas maltophilia* |
| P01324 | *Cronobacter sakazakii*, *Enterobacter cloacae*, *Enterobacteria phage HK97*, *Enterococcus casseliflavus*, *Enterococcus faecium*, *Escherichia coli*, *Klebsiella pneumoniae*, *Kocuria rhizophila*, *Lactococcus lactis*, *Leuconostoc mesenteroides*, *Micrococcus luteus*, *Pseudomonas stutzeri*, *Rhodopseudomonas palustris*, *Stenotrophomonas maltophilia*, *Stenotrophomonas phage phiSMA7*, *Streptococcus parauberis*, *Streptococcus suis*, *Streptococcus thermophilus* |

**Supplementary Table 5:**
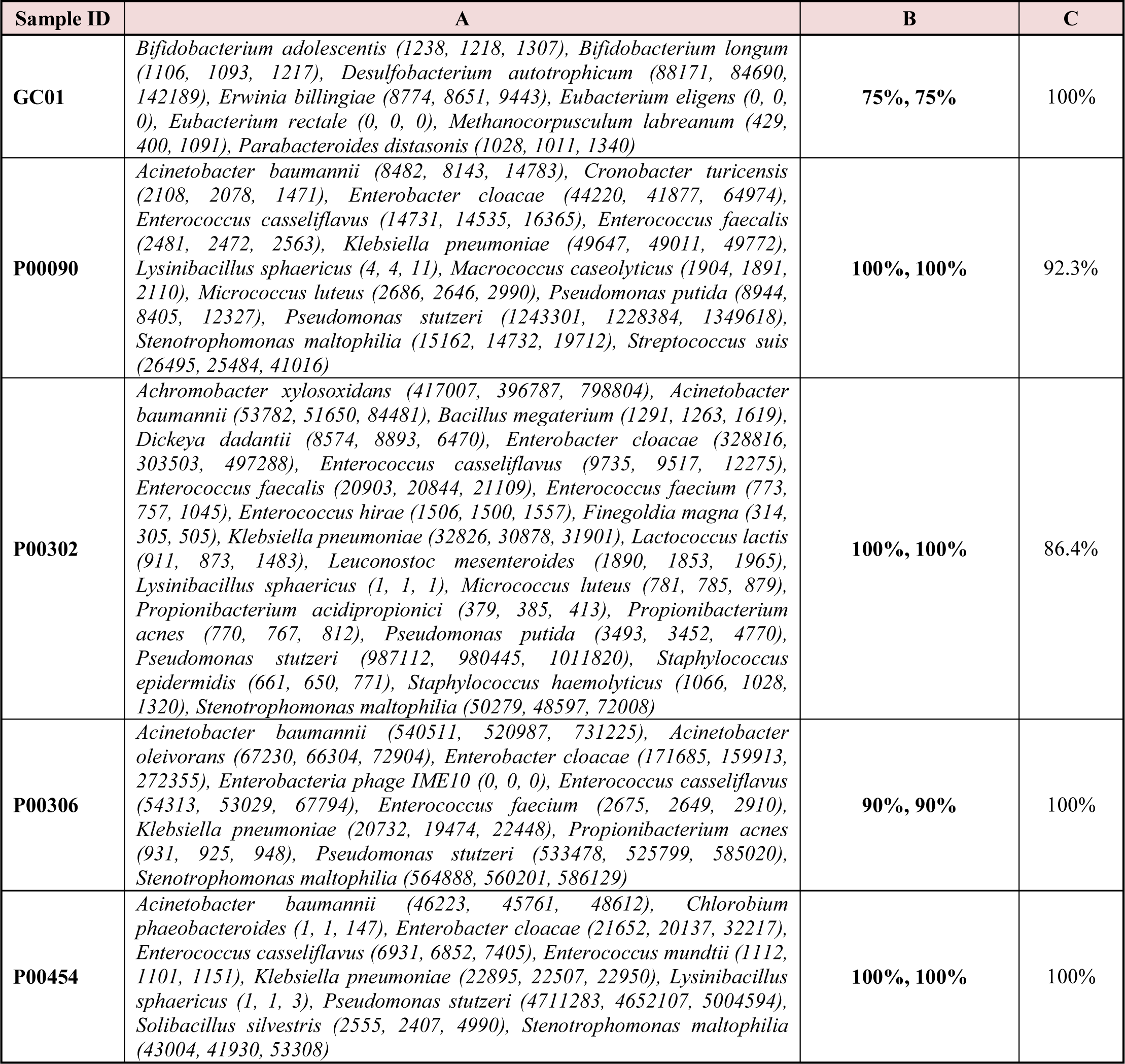
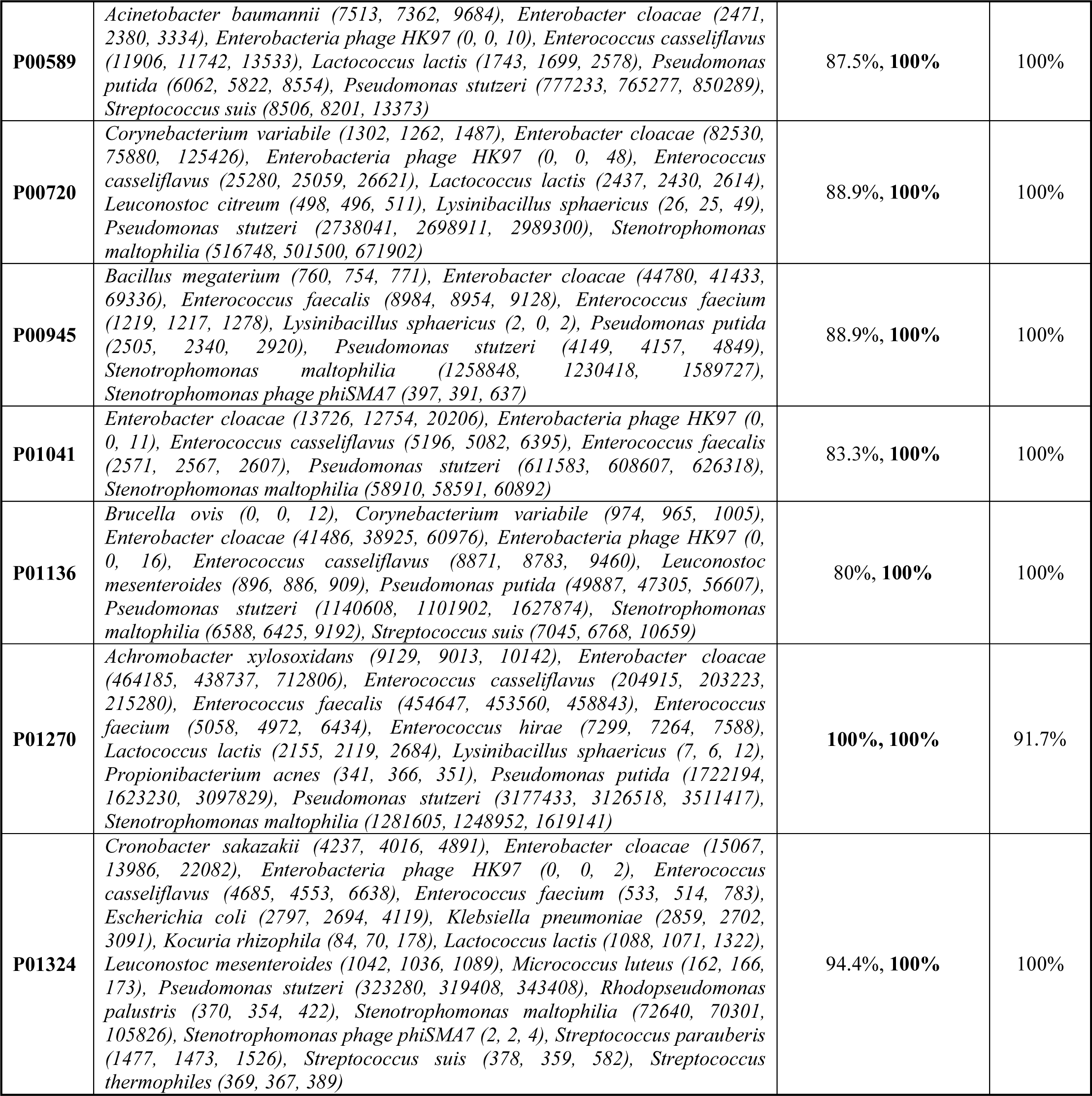
Column A lists the reads count reported by CLARK, and CLARK-S on the species listed in Supplementary Table 4. For each species, the reads count is reported as a pair (CLARK, CLARK-S). Column B reports the agreement rate between [2] and results reported by CLARK, and CLARK-S, in this order. For example, for the sample GC01, the agreement rate between CLARK and [2] was 75% because CLARK detected the presence of 6 species out of the 8 in [2]. Values in bold indicate the highest agreement rate. Column C reports the percentage of species for which CLARK-S reports a higher reads count than CLARK. For example, for the sample P00090, CLARK-S reports a higher number of reads count than CLARK for 12 species out of 13 (i.e., 92.3%).

| Sample ID | A | B | C |
| --- | --- | --- | --- |
| **GC01** | *Bifidobacterium adolescentis* (1238, 1218, 1307), *Bifidobacterium longum* (1106, 1093, 1217), *Desulfobacterium autotrophicum* (88171, 84690, 142189), *Erwinia billingiae* (8774, 8651, 9443), *Eubacterium eligens* (0, 0, 0), *Eubacterium rectale* (0, 0, 0), *Methanocorpusculum labreanum* (429, 400, 1091), *Parabacteroides distasonis* (1028, 1011, 1340) | **75%, 75%** | 100% |
| **P00090** | *Acinetobacter baumannii* (8482, 8143, 14783), *Cronobacter turicensis* (2108, 2078, 1471), *Enterobacter cloacae* (44220, 41877, 64974), *Enterococcus casseliflavus* (14731, 14535, 16365), *Enterococcus faecalis* (2481, 2472, 2563), *Klebsiella pneumoniae* (49647, 49011, 49772), *Lysinibacillus sphaericus* (4, 4, 11), *Macrococcus caseolyticus* (1904, 1891, 2110), *Micrococcus luteus* (2686, 2646, 2990), *Pseudomonas putida* (8944, 8405, 12327), *Pseudomonas stutzeri* (1243301, 1228384, 1349618), *Stenotrophomonas maltophilia* (15162, 14732, 19712), *Streptococcus suis* (26495, 25484, 41016) | **100%, 100%** | 92.3% |
| **P00302** | *Achromobacter xylosoxidans* (417007, 396787, 798804), *Acinetobacter baumannii* (53782, 51650, 84481), *Bacillus megaterium* (1291, 1263, 1619), *Dickeya dadantii* (8574, 8893, 6470), *Enterobacter cloacae* (328816, 303503, 497288), *Enterococcus casseliflavus* (9735, 9517, 12275), *Enterococcus faecalis* (20903, 20844, 21109), *Enterococcus faecium* (773, 757, 1045), *Enterococcus hirae* (1506, 1500, 1557), *Fingoldia magna* (314, 305, 505), *Klebsiella pneumoniae* (32826, 30878, 31901), *Lactococcus lactis* (911, 873, 1483), *Leuconostoc mesenteroides* (1890, 1853, 1965), *Lysinibacillus sphaericus* (1, 1, 1), *Micrococcus luteus* (781, 785, 879), *Propionibacterium acidipropionici* (379, 385, 413), *Propionibacterium acnes* (770, 767, 812), *Pseudomonas putida* (3493, 3452, 4770), *Pseudomonas stutzeri* (987112, 980445, 1011820), *Staphylococcus epidermidis* (661, 650, 771), *Staphylococcus haemolyticus* (1066, 1028, 1320), *Stenotrophomonas maltophilia* (50279, 48597, 72008) | **100%, 100%** | 86.4% |
| **P00306** | *Acinetobacter baumannii* (540511, 520987, 731225), *Acinetobacter oleivorans* (67230, 66304, 72904), *Enterobacter cloacae* (171685, 159913, 272355), *Enterobacteria phage IME10* (0, 0, 0), *Enterococcus casseliflavus* (54313, 53029, 67794), *Enterococcus faecium* (2675, 2649, 2910), *Klebsiella pneumoniae* (20732, 19474, 22448), *Propionibacterium acnes* (931, 925, 948), *Pseudomonas stutzeri* (533478, 525799, 585020), *Stenotrophomonas maltophilia* (564888, 560201, 586129) | **90%, 90%** | 100% |
| **P00454** | *Acinetobacter baumannii* (46223, 45761, 48612), *Chlorobium phaeobacteroides* (1, 1, 147), *Enterobacter cloacae* (21652, 20137, 32217), *Enterococcus casseliflavus* (6931, 6852, 7405), *Enterococcus mundtii* (1112, 1101, 1151), *Klebsiella pneumoniae* (22895, 22507, 22950), *Lysinibacillus sphaericus* (1, 1, 3), *Pseudomonas stutzeri* (4711283, 4652107, 5004594), *Solibacillus silvestris* (2555, 2407, 4990), *Stenotrophomonas maltophilia* (43004, 41930, 53308) | **100%, 100%** | 100% |
| **P00589** | *Acinetobacter baumannii* (7513, 7362, 9684), *Enterobacter cloacae* (2471, 2380, 3334), *Enterobacteria phage HK97* (0, 0, 10), *Enterococcus casseliflavus* (11906, 11742, 13533), *Lactococcus lactis* (1743, 1699, 2578), *Pseudomonas putida* (6062, 5822, 8554), *Pseudomonas stutzeri* (777233, 765277, 850289), *Streptococcus suis* (8506, 8201, 13373) | 87.5%, **100%** | 100% |
| **P00720** | *Corynebacterium variabile* (1302, 1262, 1487), *Enterobacter cloacae* (82530, 75880, 125426), *Enterobacteria phage HK97* (0, 0, 48), *Enterococcus casseliflavus* (25280, 25059, 26621), *Lactococcus lactis* (2437, 2430, 2614), *Leuconostoc citreum* (498, 496, 511), *Lysinibacillus sphaericus* (26, 25, 49), *Pseudomonas stutzeri* (2738041, 2698911, 2989300), *Stenotrophomonas maltophilia* (516748, 501500, 671902) | 88.9%, **100%** | 100% |
| **P00945** | *Bacillus megaterium* (760, 754, 771), *Enterobacter cloacae* (44780, 41433, 69336), *Enterococcus faecalis* (8984, 8954, 9128), *Enterococcus faecium* (1219, 1217, 1278), *Lysinibacillus sphaericus* (2, 0, 2), *Pseudomonas putida* (2505, 2340, 2920), *Pseudomonas stutzeri* (4149, 4157, 4849), *Stenotrophomonas maltophilia* (1258848, 1230418, 1589727), *Stenotrophomonas phage phiSMA7* (397, 391, 637) | 88.9%, **100%** | 100% |
| **P01041** | *Enterobacter cloacae* (13726, 12754, 20206), *Enterobacteria phage HK97* (0, 0, 11), *Enterococcus casseliflavus* (5196, 5082, 6395), *Enterococcus faecalis* (2571, 2567, 2607), *Pseudomonas stutzeri* (611583, 608607, 626318), *Stenotrophomonas maltophilia* (58910, 58591, 60892) | 83.3%, **100%** | 100% |
| **P01136** | *Brucella ovis* (0, 0, 12), *Corynebacterium variabile* (974, 965, 1005), *Enterobacter cloacae* (41486, 38925, 60976), *Enterobacteria phage HK97* (0, 0, 16), *Enterococcus casseliflavus* (8871, 8783, 9460), *Leuconostoc mesenteroides* (896, 886, 909), *Pseudomonas putida* (49887, 47305, 56607), *Pseudomonas stutzeri* (1140608, 1101902, 1627874), *Stenotrophomonas maltophilia* (6588, 6425, 9192), *Streptococcus suis* (7045, 6768, 10659) | 80%, **100%** | 100% |
| **P01270** | *Achromobacter xylosoxidans* (9129, 9013, 10142), *Enterobacter cloacae* (464185, 438737, 712806), *Enterococcus casseliflavus* (204915, 203223, 215280), *Enterococcus faecalis* (454647, 453560, 458843), *Enterococcus faecium* (5058, 4972, 6434), *Enterococcus hirae* (7299, 7264, 7588), *Lactococcus lactis* (2155, 2119, 2684), *Lysinibacillus sphaericus* (7, 6, 12), *Propionibacterium acnes* (341, 366, 351), *Pseudomonas putida* (1722194, 1623230, 3097829), *Pseudomonas stutzeri* (3177433, 3126518, 3511417), *Stenotrophomonas maltophilia* (1281605, 1248952, 1619141) | **100%, 100%** | 91.7% |
| **P01324** | *Cronobacter sakazakii* (4237, 4016, 4891), *Enterobacter cloacae* (15067, 13986, 22082), *Enterobacteria phage HK97* (0, 0, 2), *Enterococcus casseliflavus* (4685, 4553, 6638), *Enterococcus faecium* (533, 514, 783), *Escherichia coli* (2797, 2694, 4119), *Klebsiella pneumoniae* (2859, 2702, 3091), *Kocuria rhizophila* (84, 70, 178), *Lactococcus lactis* (1088, 1071, 1322), *Leuconostoc mesenteroides* (1042, 1036, 1089), *Micrococcus luteus* (162, 166, 173), *Pseudomonas stutzeri* (323280, 319408, 343408), *Rhodopseudomonas palustris* (370, 354, 422), *Stenotrophomonas maltophilia* (72640, 70301, 105826), *Stenotrophomonas phage phiSMA7* (2, 2, 4), *Streptococcus parauberis* (1477, 1473, 1526), *Streptococcus suis* (378, 359, 582), *Streptococcus thermophiles* (369, 367, 389) | 94.4%, **100%** | 100% |

**Supplementary Table 6:** Assignment rate (i.e., ratio in percent between the number of assigned/classified reads and the total number of reads) on real samples for CLARK and CLARK-*S*. Values in bold are the highest.

| *Sample ID* | *CLARK* | *CLARK-S* |
| --- | --- | --- |
| GC01 | 1.36% | **2.55%** |
| P00090 | 49.59% | **56.16%** |
| P00302 | 23.70% | **29.89%** |
| P00306 | 33.82% | **40.47%** |
| P00454 | 66.37% | **71.50%** |
| P00589 | 29.46% | **34.24%** |
| P00720 | 55.59% | **64.35%** |
| P00945 | 23.21% | **35.65%** |
| P01041 | 50.28% | **64.35%** |
| P01136 | 26.36% | **35.65%** |
| P01270 | 50.28% | **64.35%** |
| P01324 | 23.29% | **27.23%** |

**Supplementary Table 7:** Classification speed of CLARK and CLARK-*S* on the synthetic datasets (default and unambiguous), the negative control samples and the real samples. The values are in thousand of read per minute. Values in bold are the highest.

| Default | *CLARK (1 CPU)* | *CLARK-S (1 CPU)* | *CLARK-S (8 CPUs)* |
| --- | --- | --- | --- |
| Buc12 | **4, 839.5** | 214.4 | 1, 220.8 |
| CParMed48 | **3, 691.4** | 204.3 | 913.6 |
| Gut20 | **3, 369.5** | 196.1 | 1, 077.8 |
| Hou31 | **3, 465.5** | 201.4 | 1, 067.7 |
| Hou21 | **3, 308.9** | 199.2 | 1, 124.6 |
| Soi50 | **3, 193.3** | 169.5 | 1, 074.7 |
| simBA-525 | **3, 194.5** | 203.1 | 1, 092.5 |
| Unambiguous | *CLARK (1 CPU)* | *CLARK-S (1 CPU)* | *CLARK-S (8 CPUs)* |
| Buc12 | **4, 160.5** | 217.7 | 1, 101.5 |
| CParMed48 | **4, 057.7** | 201.3 | 874.1 |
| Gut20 | **2, 954.0** | 134.3 | 1, 083.7 |
| Hou31 | **3, 912.9** | 142.0 | 964.0 |
| Hou21 | **3, 801.1** | 157.8 | 1, 003.8 |
| Soi50 | **2, 868.9** | 141.4 | 1, 024.7 |
| simBA-525 | **3, 359.0** | 141.7 | 1, 076.3 |

| Negative control | *CLARK (1 CPU)* | *CLARK-S (1 CPU)* | *CLARK-S (8 CPUs)* |
| --- | --- | --- | --- |
| HM1 | **2, 619.1** | 146.2 | 1, 033.1 |
| HM2 | **2, 932.1** | 131.9 | 937.9 |
| LM | **2, 654.2** | 134.2 | 957.3 |

| Sample ID | *CLARK (1 CPU)* | *CLARK-S (1 CPU)* | *CLARK-S (8 CPUs)* |
| --- | --- | --- | --- |
| GC01 | **3,142.3** | 290.7 | 1,315.9 |
| P00090 | **2,587.7** | 230.7 | 1,355.7 |
| P00302 | **3,330.3** | 326.7 | 1,432.1 |
| P00306 | **3,553.6** | 332.5 | 1,428.1 |
| P00454 | **3,668.7** | 364.7 | 1,569.5 |
| P00589 | **4,929.9** | 312.2 | 1,373.8 |
| P00720 | **5,203.0** | 312.2 | 1,545.8 |
| P00945 | **4,758.7** | 324.2 | 1,390.9 |
| P01041 | **4,348.5** | 313.9 | 1,381.2 |
| P01136 | **4,893.1** | 315.0 | 1,371.2 |
| P01270 | **3,548.8** | 341.8 | 1,531.8 |
| P01324 | **3,513.6** | 320.1 | 1,363.9 |

**Supplementary Table 8:**
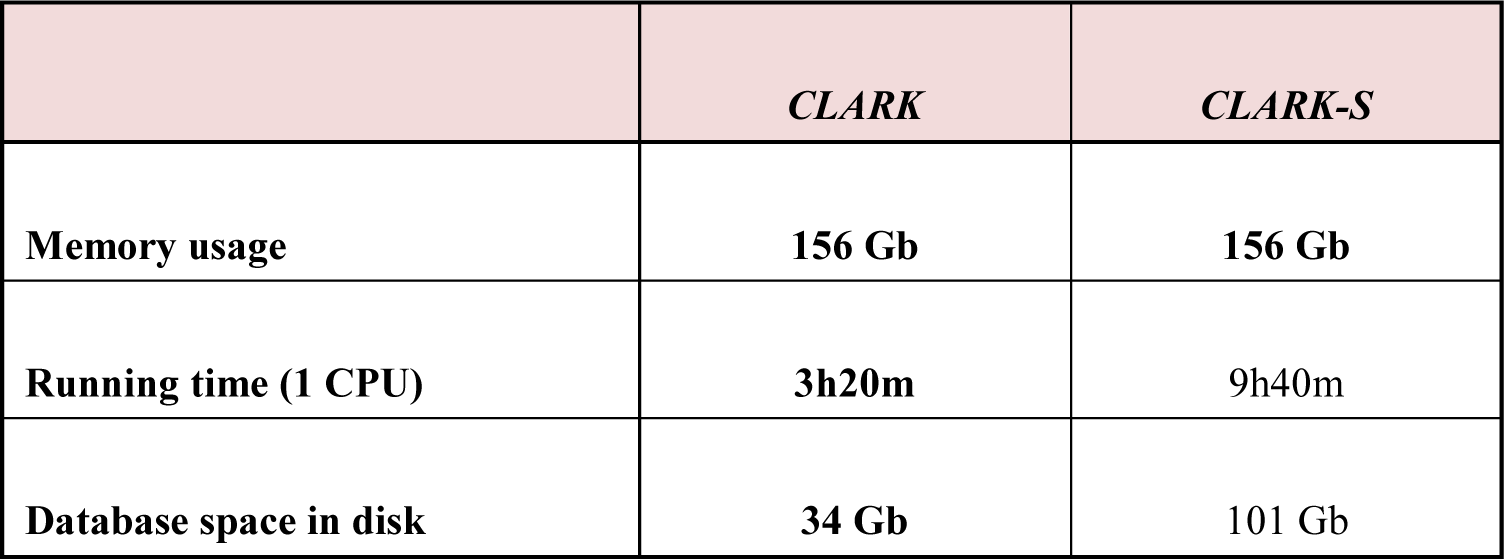
Memory usage and running time for the index creation for CLARK and CLARK-S. The database is the bacterial, archaeal and viral sequences from NCBI/RefSeq. Measures indicated were obtained via the “/usr/bin/time–v” command. All tools CLARK and CLARK-S (v1.2.2-b) were run on a Linux server (20 cores Intel Xeon CPU E5-2690v2 3.3GHz and 512GB of RAM). Lowest values are indicated in bold.

|  | *CLARK* | *CLARK-S* |
| --- | --- | --- |
| Memory usage | **156 Gb** | **156 Gb** |
| Running time (1 CPU) | **3h20m** | 9h40m |
| Database space in disk | **34 Gb** | 101 Gb |

**Supplementary Figure 1:**
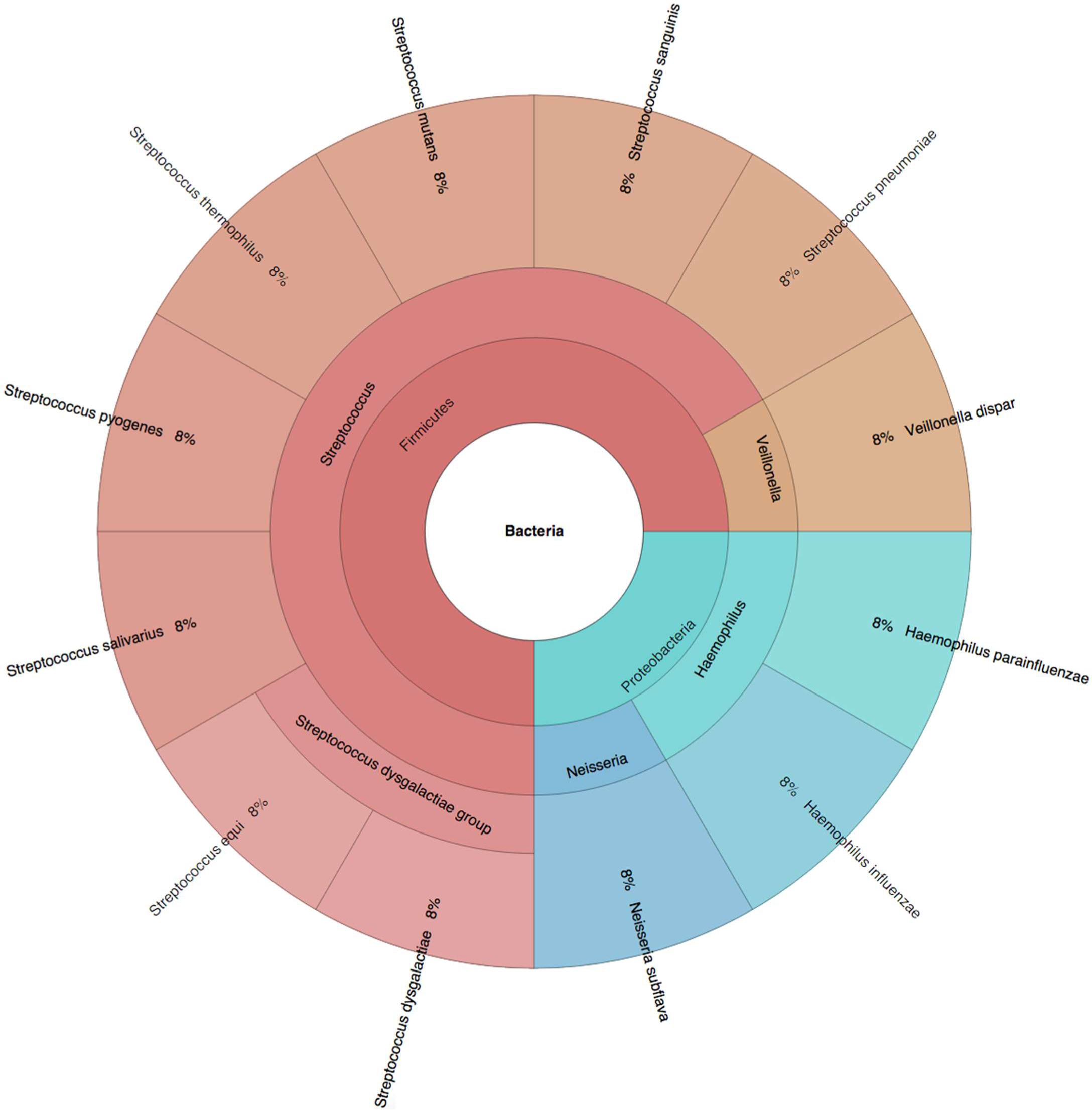
Buc12.

**Supplementary Figure 2:**
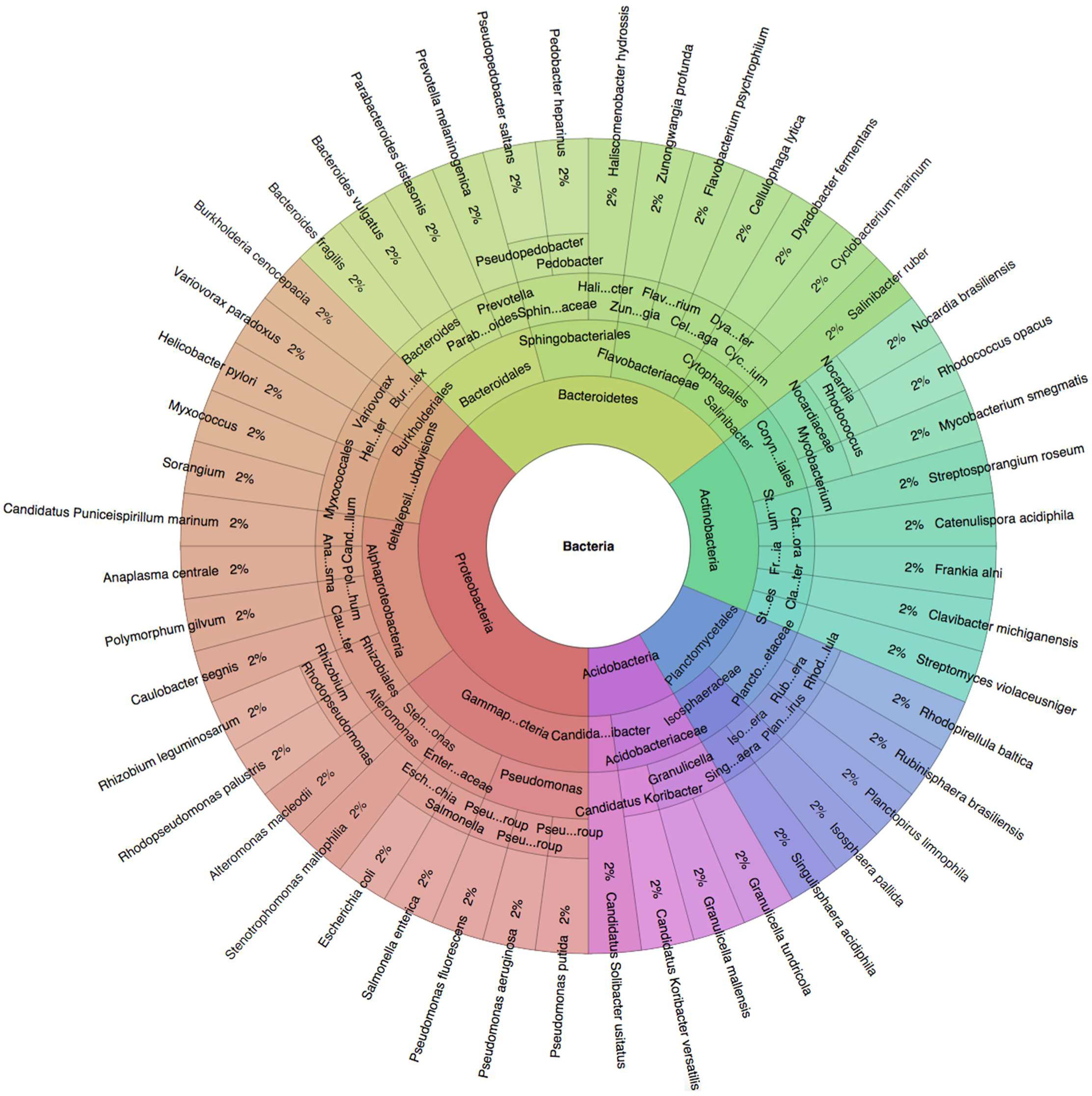
CParMed48.

**Supplementary Figure 3:**
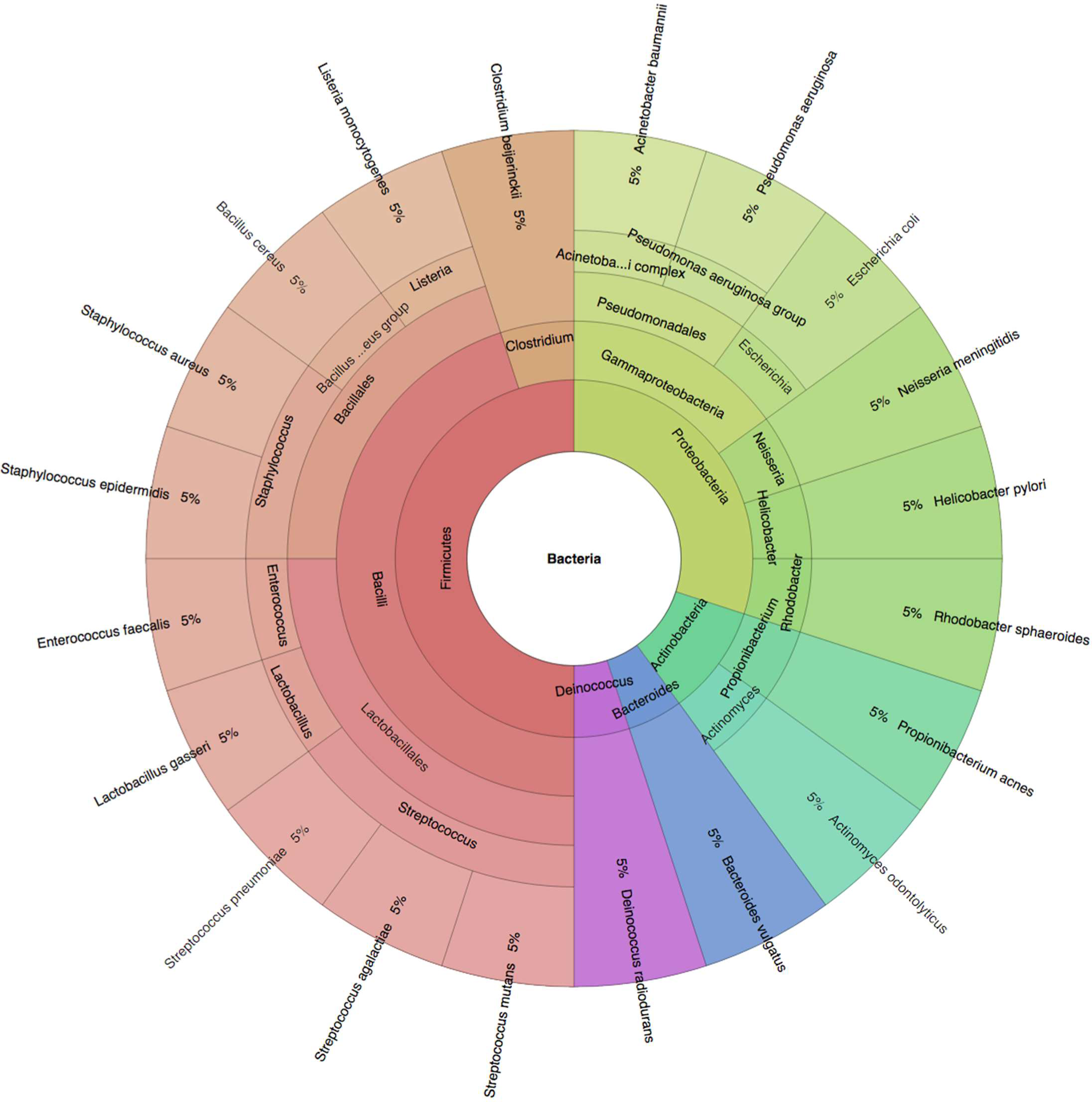
Gut20.

**Supplementary Figure 4:**
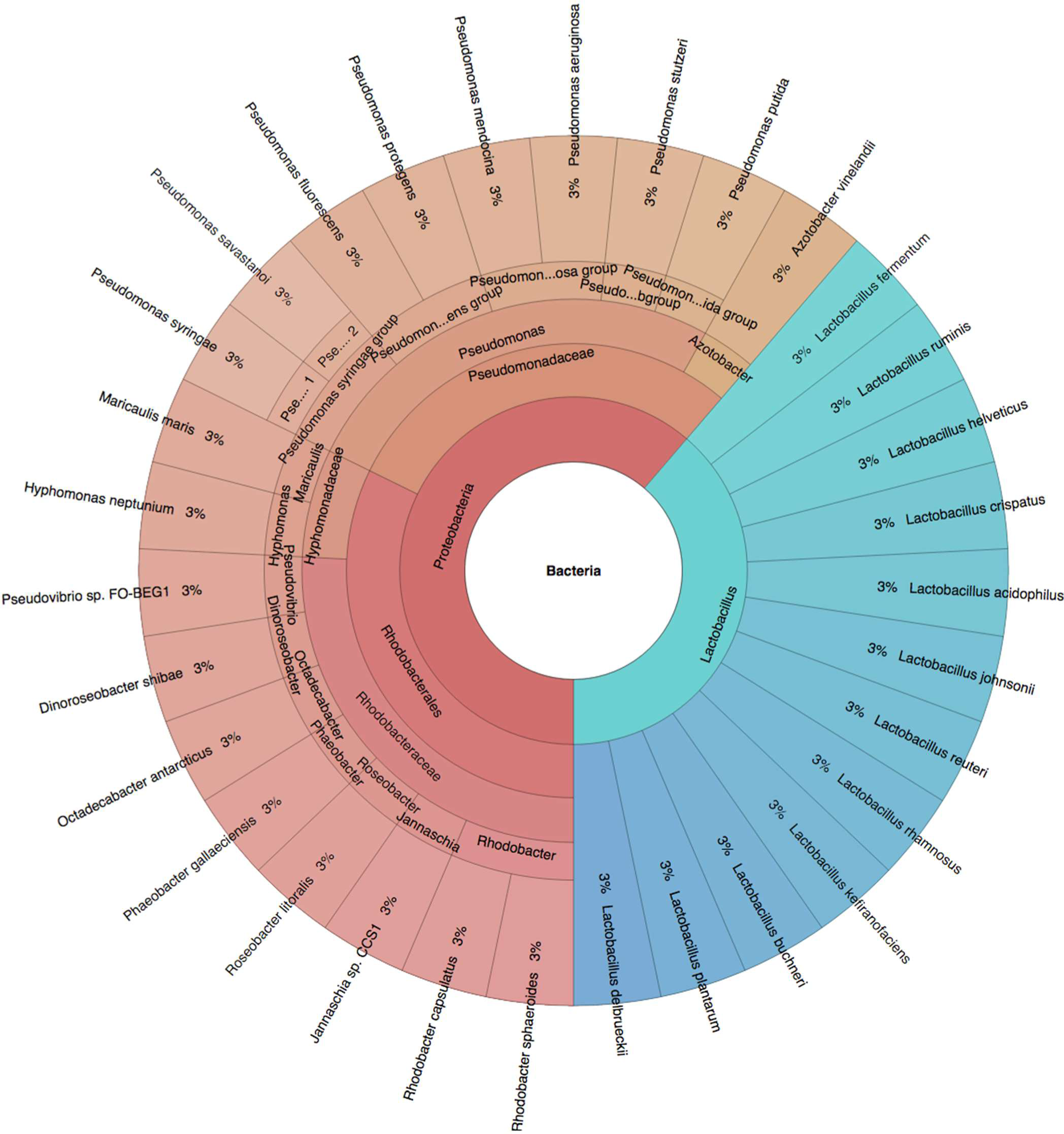
Hous31.

**Supplementary Figure 5:**
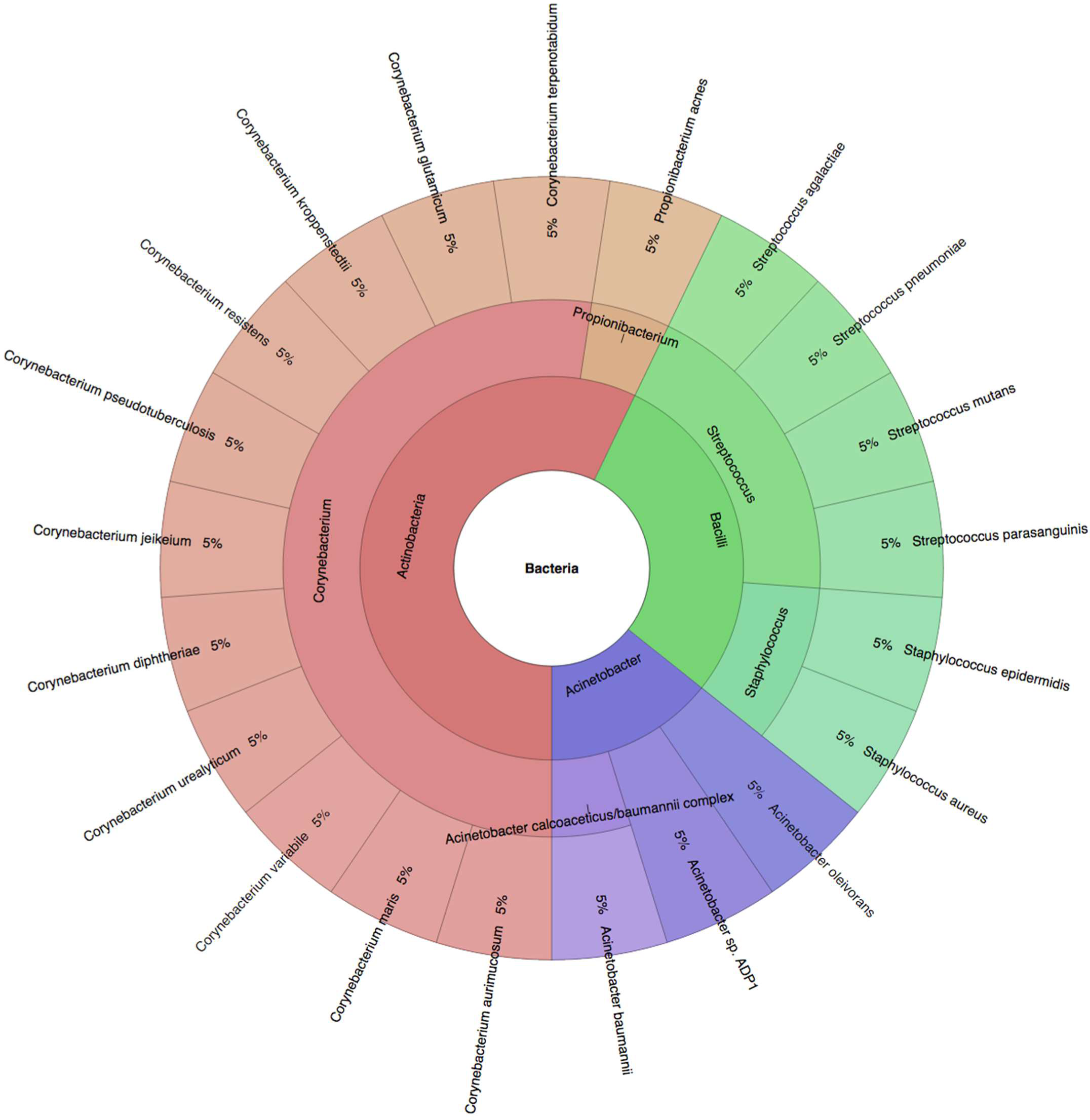
Hous21.

**Supplementary Figure 6:**
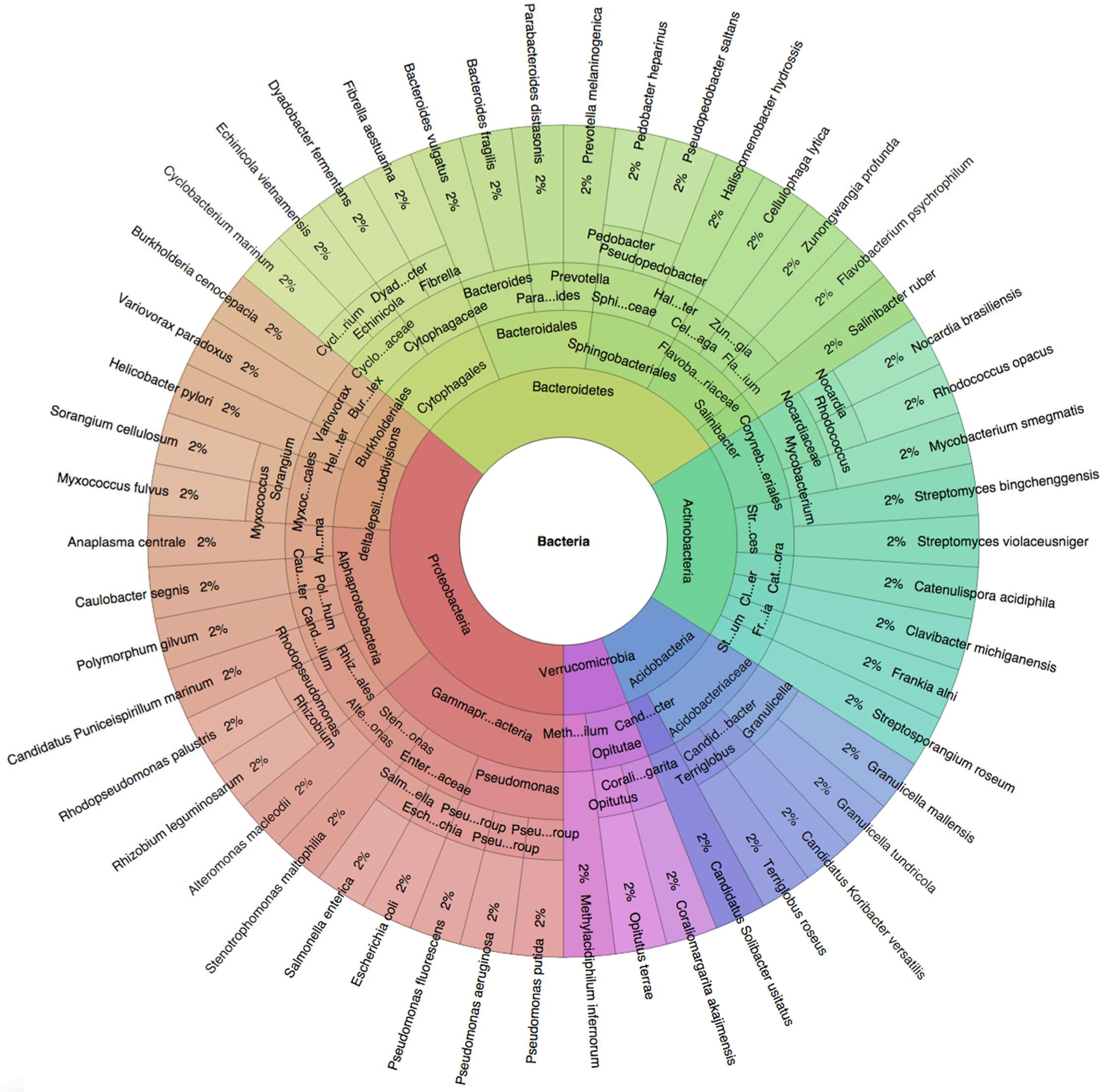
Soi50.

**Supplementary Figure 7:**
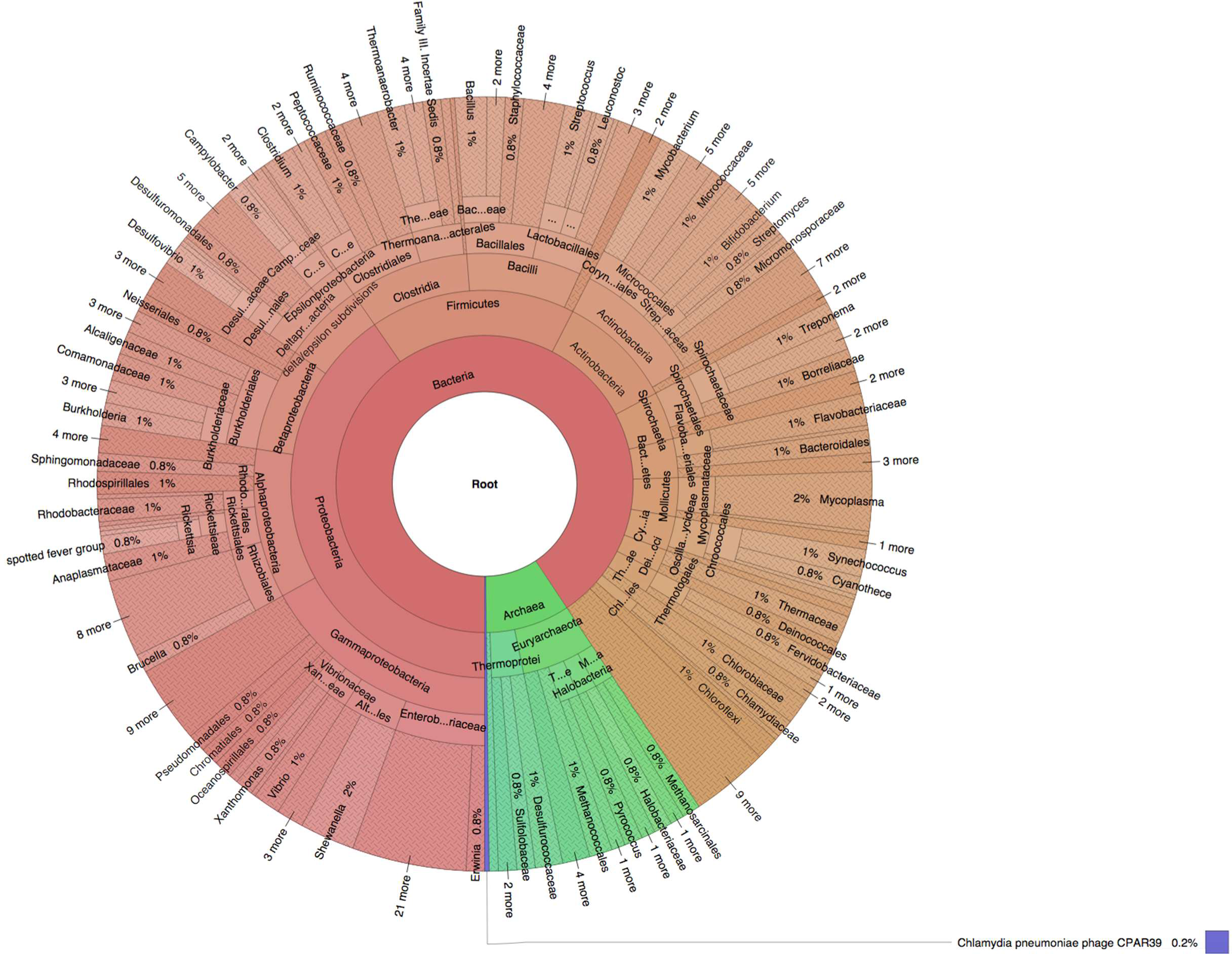
simBA-525.

## References

[1] Venter, J. C., Remington, K., Heidelberg, J. F., Halpern, A. L., Rusch, D., Eisen, J. A., Wu, D., Paulsen, I., Nelson, K. E., et al.(2004). Environmental genome shotgun sequencing of the Sargasso Sea. Science, 304(5667), 66–74.

[2] Huttenhower, C., Gevers, D., Knight, R., Abubucker, S., Badger, J., and Chinwalla, A. e. a. (2012). Structure, function and diversity of the healthy human microbiome. Nature, 486(7402), 207–214.

[3] Ounit, R., Wanamaker, S., Close, T. J., Lonardi, S. (2015). CLARK: fast and accurate classification of metagenomic and genomic sequences using discriminative k-mers. BMC Genomics, 16(1), 236.

[4] Lindgreen, S., Adair, K. L., et al. (2016). An evaluation of the accuracy and speed of metagenome analysis tools. Scientific reports, 6, 19233.

[5] Ma, B., Tromp, J., et al. (2002). PatternHunter: faster and more sensitive homology search. Bioinformatics, 18(3), 440–445.

[6] Huson, D. H., Auch, A. F., et al. (2007). MEGAN analysis of metagenomic data. Genome Research, 17(3), 377–386.

[7] Brown, D. G., Li, M., et al. (2004). A tutorial of recent developments in the seeding of local alignment. Journal of Bioinformatics and Computational Biology, 2(04), 819–842.

[8] Ilie, L., Ilie, S., et al. (2011). SpEED: fast computation of sensitive spaced seeds. Bioinformatics, 27(17), 2433–2434.

[9] Huang, W., Li, L., et al. (2012). ART: a next-generation sequencing read simulator. Bioinformatics, 28(4), 593–594.

[10] Afshinnekoo, E., Meydan, C., et al. (2015). Geospatial resolution of human and bacterial diversity with city-scale metagenomics. Cell Systems, 1(1), 72–87.

[11] Thompson, L. R., Williams, G. J., et al. (2016). Metagenomic covariation along densely sampled environmental gradients in the red sea. The ISME journal.

## References

[1] Adams, R. I., Bateman, A. C., Bik, H. M., and Meadow, J. F. Microbiota of the indoor environment: a meta-analysis. Microbiome 3, 1 (2015), 1.

[2] Afshinnekoo, E., Meydan, C., Chowdhury, S., Jaroudi, D., Boyer, C., Bernstein, N., Maritz, J. M., Reeves, D., Gandara, J., Chhangawala, S., et al. Geospatial resolution of human and bacterial diversity with city-scale metagenomics. Cell systems 1, 1 (2015), 72–87.

[3] Fierer, N., Leff, J. W., Adams, B. J., Nielsen, U. N., Bates, S. T., Lauber, C. L., Owens, S., Gilbert, J. A., Wall, D. H., and Caporaso, J. G. Cross-biome metagenomic analyses of soil microbial communities and their functional attributes. Proceedings of the National Academy of Sciences 109, 52 (2012), 21390–21395.

[4] Franzosa, E. A., Huang, K., Meadow, J. F., Gevers, D., Lemon, K. P., Bohannan, B. J., and Huttenhower, C. Identifying personal microbiomes using metagenomic codes. Proceedings of the National Academy of Sciences 112, 22 (2015), E2930–E2938.

[5] Huttenhower, C., Gevers, D., Knight, R., Abubucker, S., Badger, J., and Chinwalla, A. e. a. Structure, function and diversity of the healthy human microbiome. Nature 486, 7402 (2012), 207–214.

[6] Huang, W., Li, L., Myers, J. R., and Marth, G. T. Art: a next-generation sequencing read simulator. Bioinformatics 28, 4 (2012), 593–594.

[7] Kuleshov, V., Jiang, C., Zhou, W., Jahanbani, F., Batzoglou, S., and Snyder, M. Synthetic long-read sequencing reveals intraspecies diversity in the human microbiome. Nature biotechnology 34, 1 (2016), 64–69.

[8] Ondov, B. D., Bergman, N. H., and Phillippy, A. M. (2011). Interactive metagenomic visualization in a web browser. BMC bioinformatics, 12(1), 1.

[9] Reese, A. T., Savage, A., Youngsteadt, E., McGuire, K. L., Koling, A., Watkins, O., Frank, S. D., and Dunn, R. R. Urban stress is associated with variation in microbial species composition - but not richness - in manhattan. The ISME journal(2015).

[10] Ruiz-Calderon, J. F., Cavallin, H., Song, S. J., Novoselac, A., Pericchi, L. R., Hernandez, J. N., Rios, R., Branch, O. H., Pereira, H., Paulino, L. C., et al. Walls talk: Microbial biogeography of homes spanning urbanization. Science advances 2, 2 (2016), e1501061.

